# Analysis of a fully infectious bioorthogonally modified human virus reveals novel features of virus cell entry

**DOI:** 10.1101/693036

**Authors:** Remigiusz A. Serwa, Eiki Sekine, Jonathan Brown, Su Hui Catherine Teo, Edward W. Tate, Peter O’Hare

## Abstract

We report the analysis of a complex enveloped human virus, herpes simplex virus (HSV), assembled after in vivo incorporation of bio-orthogonal methionine analogues homopropargylglycine (HPG) or azidohomoalanine (AHA). We optimised protocols for the production of virions incorporating AHA (termed HSV^AHA^), identifying conditions which resulted in normal yields of HSV and normal particle/pfu ratios. Moreover we show that essentially every single HSV^AHA^ capsid-containing particle was detectable at the individual particle level by chemical ligation of azide-linked fluorochromes to AHA-containing structural proteins. This was a completely specific chemical ligation, with no capsids assembled under normal methionine-containing conditions detected in parallel. We demonstrate by quantitative mass spectrometric analysis that HSV^AHA^ virions exhibit no qualitative or quantitative differences in the repertoires of structural proteins compared to virions assembled under normal conditions. Individual proteins and AHA incorporation sites were identified in capsid, tegument and envelope compartments, including major essential structural proteins. Finally we revealing novel aspects of entry pathways using HSV^AHA^ and chemical fluorochrome ligation that were not apparent from conventional immunofluorescence. Since ligation targets total AHA- containing protein and peptides, our results demonstrate the presence of abundant AHA-labelled products in cytoplasmic macrodomains and tubules which no longer contain intact particles detectable by immunofluorescence. Although these do not co-localise with lysosomal markers, we propose they may represent sites of proteolytic virion processing. Analysis of HSV^AHA^ also enabled the discrimination or primary entering from secondary assembling, demonstrating assembly and second round infection within 6 hrs of initial infection and dual infections of primary and secondary virus in spatially restricted cytoplasmic areas of the same cell. Together with other demonstrated applications e.g., in genome biology, lipid and protein trafficking, the work further exemplifies the utility and potential of bio-orthogonal chemistry for studies in many aspects of virus-host interactions.

**Author Summary:** Bio-orthogonal chemistry and the application of non-canonical amino acid tagging (BONCAT) has opened opportunities for analysis of translational control, protein trafficking and modification in studies of infection and immunity. We expand on our earlier work, reporting the tractable, scalable production and analysis of a large structurally complex enveloped human virus that incorporates non-canonical amino acids into structural proteins from all parts of the virus particle. Thus in the complex translationally altered environment of a herpesvirus infected cell, non-canonical amino acid incorporation has no significant functional effect on the multiple cellular and viral functions required to assemble infectious virions. We further demonstrate the normal recruitment and stochiometries of all detected structural proteins, which combined with data showing unaltered virus yields and infectivity, indicates that the non-canonical residues in the various structural proteins have no effect on their subsequent ability to infect cells. Focusing on spatial analysis of virus entry, we provide examples and reveal novel insight into virus entry and processing. Together with previous reports using bio-orthogonal chemistry in studies of virus infection the potential applications of BMVs are considerable.

## Introduction

Advances in the field of biological chemistry, particularly the development of bio-orthogonal non-canonical amino acid tagging (BONCAT), have opened new opportunities for analysis of protein synthesis and modification in the field of microbial infection and immunity. Azido-bearing AHA or alkyne-bearing HPG are analogues of methionine containing small chemical moieties that have no overall charge and are well tolerated for protein incorporation in vivo [1, 2]. AHA or HPG incorporation combined with CuAAC covalent ligation to reciprocal fluorescent capture agents [3, 4] enables spatial and biochemical analysis of newly translated proteins [5–10]. This can be further combined with selective protein isolation offering facilitating integrated spatiotemporal, biochemical and systems characterisation of newly translated proteins during infection. We previously used bio-orthogonal labelling to investigate lipidation, protein trafficking and genome entry during herpes simplex virus infection [11–14] yielding new insight into HSV genome biology and translational control. Although we did not extend the work into examination of HSV assembly and virion protein modification in the presence of bio-orthogonal amino acids, several previous reports have demonstrated the potential of chemical biology to investigate virus or virus-like particles [15, 16]. In a simplified system using bacterial expression in AHA-containing medium of individual proteins of hepatitis B virus (HBV) or bacteriophage Qβ, it was reported that coat proteins incorporating AHA could assemble into virus-like particles. While this was not a study to make complex infectious bio-orthogonally labelled viruses in mammalian culture systems, it demonstrated the principle of virus particle assembly with bio-orthogonal amino acids, albeit from a single protein in a simplified system.

Another approach combined genetic engineering with bio-orthogonal chemistry [16, 17]. Here an amber stop codon is first introduced into one specific individual target protein (HIV envelope protein in this case), by site-directed mutagenesis. Upon co-expression of a separate aminoacyl tRNA synthase/tRNA pair, a non-canonical amino acid can be incorporated into the amber codon of the mutated gene, resulting in translation through the stop codon. This approach while useful has drawbacks, focuses on a single viral protein and also requires a complex experimental set-up, including constructs to express the amber stop codon, a pyrrolysyl-tRNA synthase and a tRNA (tRNApyl/pylRSAF) for amber codon recognition, and a mutated release factor to augment amber suppression, which was nevertheless incomplete [16]. Yet another approach reported chemical labelling of vaccinia virus with Quantum Dots (QDs) [18]. In this case however, the virus was produced under normal conditions without any bio-orthogonal precursors and then had to be modified non-specifically by *N*-hydroxysuccinimidyl (NHS) ester coupling of an azide moiety to general amine groups on surface proteins and, in a second step, then ligated to QDs that had been modified with dibenzocyclooctynes (DBCO). Not only is this also a complex pipeline, general NHS-ester modification of amine groups on virus surface proteins is likely to have a detrimental effect on infectivity and imaging data showed that modified particles could only be detected on the outside of cells.

One previous report has indicated the possibility of in vivo labelling with bio-orthogonal amino acid precursors for the production of adenovirus particles incorporating AHA in structural proteins without loss of Ad particle infectivity [19]. In this work, adenovirus type 5 (Ad5) infected 293T cells were pulsed with AHA containing medium for 6 hrs and then chased in normal methionine containing medium for a further 24 hrs. AHA-labelled Ad5 particles were recovered and AHA incorporation into a number of structural proteins reported from mass spectrometry. The efficiency of detection of individual particles by click-ligation to structural proteins coupled with imaging analysis was not assessed. However results from click ligation of a fluorochrome on the intact Ad5 virions, followed by SDS-PAGE of disrupted particles, indicated that 5-10% of exposed methionines could be coupled by CuAAC ligation. Although the comparative effect of AHA labelling compared to normal virus production in culture on the total repertoire of Ad5 proteins was not reported, the authors reported that the virus preparations retained full infectivity. They further used the approach to tag particles with an alkyne-PEGylated-folate moiety and reported considerably increased infection in a folate-receptor bearing cell line.

Here we develop a comparatively simple and tractable approach for the production and investigation of a complex bio-orthogonally modified enveloped virus HSV. We show such virions, which we generally term BMVs (bio-orthogonally modified viruses) are assembled normally, are fully infectious and importantly retain the normal repertoire of structural proteins as assessed by quantitative mass spectrometry compared to normal virions assembled in methionine containing medium. Such viruses offer novel opportunities for the investigation of virus infection. As an example we undertake a spatiotemporal investigation of virus particles and the fate of the total protein constituents during cell entry. We undertake simultaneous analysis using click ligation of an alkyne-fluorochrome versus conventional immunofluorescence which normally uses protein/epitope-specific antibodies. The results reveal new aspects of particle cytoplasmic processing and allows spatial discrimination of infecting versus progeny viruses which is not possible with conventional immunofluorescence and add to the expanding opportunities for bio-orthogonal chemistry in studies of infection and immunity.

## Results

### Bio-orthogonal amino acid incorporation during HSV replication

During HSV infection, cellular protein synthesis is progressively suppressed and at later times mainly directed to the production of viral proteins, especially the abundant structural proteins that compose virion particles [20, 21]. Since we aimed to produce progeny virus incorporating bio-orthogonal amino acids in structural proteins, we first examined protein synthesis for extended times up to 20 hrs, the general duration of a replication cycle. Cells were infected or mock-infected as described in methods and labelled with 1 mM AHA (60 min duration) at intervals up to 20 hrs post infection (hpi). Proteins were then analysed by total protein staining (Fig 1a, lanes 1-6) or by CuAAC ligation to the fluorescent probe alkynyl-TAMRA-Biotin (YnTB) [22] and in-gel fluorescence to visualise de novo synthesised proteins (Fig 1a, lanes 7-12). The total abundance of proteins changed little although abundant viral proteins, such as VP5, could be observed (Fig 1a, VP5 arrowed). For de novo protein synthesis, as expected a number of novel species were observed by 2-4 hrs with accumulation of distinct new bands at later times and final overall decline in active protein synthesis (Fig 1a, lanes 8-12, arrowed).

**Fig 1.**
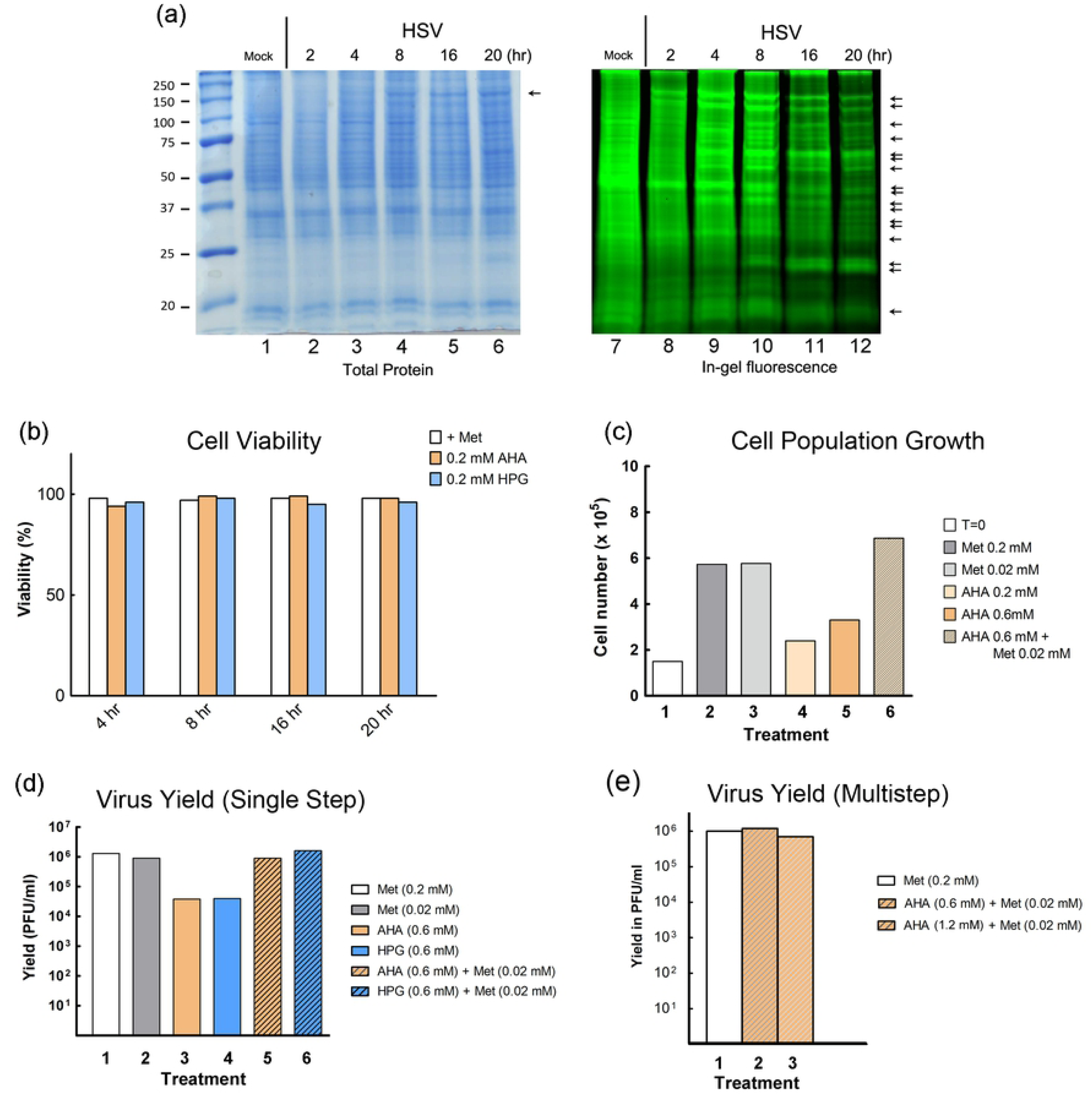
Analysis of newly synthesised proteins, cell viability, growth and virus yield in the presence of noncanonical methionine analogues. (a) Mock-infected or HSV-infected cells (moi 10) cells were pulse-labelled as described in the text. Proteins were visualised after SDS-PAGE by total protein staining or in-gel fluorescence. (b) Cells were incubated in duplicate without or with 0.1 mM AHA or HPG for durations indicated and the percentage viability assayed (viability in normal medium was set to 100%). Data is plotted as the mean of the duplicate samples. (c) Cells were plated in triplicate at a density of 1 × 10^6^ and incubated under various conditions. After 3 days, numbers of viable cells were evaluated and plotted as the mean of the triplicates for each condition. (d) Cells were infected at moi 5 and incubated under various conditions from 2-24 hpi. Extracellular virus was harvested and titres quantitated by plaque assay. (e) Virus yield after multi-round replication cycle (moi 0.005) in the presence of Met, or Met and AHA mixtures as indicated and labelled from 2-72 hpi.

### Assembly of infectious virus containing bio-orthogonal amino acids

Normal production of viruses in culture generally utilises initial low mois (0.001-0.01), so that only a small population of cells is initially infected. With the aim of efficient modified virus production, we first used standard multi-round infection after low moi infection, to generate HSV theoretically containing structural proteins incorporating either AHA or HPG (termed for ease of reference, HSV^AHA^ and HSV^HPG^). However with complete depletion of methionine from the medium and substitution with AHA or HPG, HSV plaque formation (which reflects multi-round replication) was substantially reduced. This could relate not only to host and/or virus protein synthesis but also to other pathways of amino acid metabolism. Nonetheless, consistent with previous analysis [12, 23], we found that methionine depletion with bio-orthogonal amino acid labelling did not affect overall cell viability (Fig 1b). However we did detect an effect when measuring overall cell population numbers (Fig 1c). While control cultures underwent 2-3 population doublings in the time (Fig 1c, conditions 1-3), cultures completely depleted of methionine and substituted with AHA exhibited lower population growth (Fig 1c, condition 4). Increasing AHA concentrations by 3-fold had only a marginal effect (Fig 1c, condition 5).

Thus, although AHA and HPG exhibit no detectable effect on cell viability or protein synthetic rates [12, 24], during prolonged incubation they do not completely substitute for methionine. Consistent with this, it has been reported that prolonged AHA labelling in the complete absence of methionine can result in changes in abundance of many cellular proteins [25]. However, it was observed that a small supplemental percentage of methionine in the presence of excess AHA, restored normal protein abundance, even over a 24 hrs labelling interval. We examined whether the cytostatic effects of complete depletion could be reversed by supplementing labelling medium with methionine at low concentrations. Our results showed that addition of methionine at a ratio of AHA/Met of 1/30^th^ (0.6 mM AHA/0.02 mM Met) restored cell growth rate to normal (Fig 1c, condition 6). Thus AHA was not itself detrimental and the population effect appeared to be due to the absence of methionine. Whatever the precise explanation, we next evaluated whether supplemental methionine would also restore normal viral yields. The results both for single-step replication over 22 hrs (Fig 1d, compare 3,4 with 5,6) or for multi-round replication (Fig 1e, compare conditions 1-3), showed that inclusion of 1/30^th^ the normal concentration of methionine rescued viral yields to essentially normal levels seen with methionine only.

Methionine supplementation while restoring virus yields is likely to have an effect on AHA protein incorporation, To examine this, using two strains of HSV, strains [17] and [KOS], we compared incorporation of HPG and AHA (Fig 2, H and A), under the following conditions: only H (lanes 2,7,12); only A (lanes 4,9,14); H supplemented with 1/30^th^ methionine (H/M, lanes 3,8,13); or A supplemented with 1/30^th^ methionine (A/M, lanes 5,10,15). We also examined mock-infected cells (lanes 1-5) and included controls with methionine alone (M) which showed some low level background fluorescence in certain species (asterisks, lanes 1,6,11). Efficient incorporation was observed for HPG alone (lanes 2,7,12) or AHA alone (lanes 4,9,14). As before, HSV induced a series of new species representing the major virus-encoded proteins (lanes 7 and 9; or 12 and 14, arrows). With limiting methionine supplementation (H/M or A/M) incorporation of HPG and AHA was discernibly reduced (lanes 8,10 and 13,15 respectively). This reduction is anticipated due to competition with the methionine supplement. However even under these conditions, readily detectable levels of HPG or AHA incorporation were still observed in HSV induced proteins. Taken together, these data show that inclusion of low ratios of methionine in the labelling medium restored overall yields of infectious virus at the cost of reduced but readily detectable bio-orthogonal amino acid intracellular incorporation into virus proteins. Nonetheless these data do not demonstrate that the virus proteins incorporating bio-orthogonal amino acids have been recruited into structural proteins in assembled virions.

**Fig 2.**
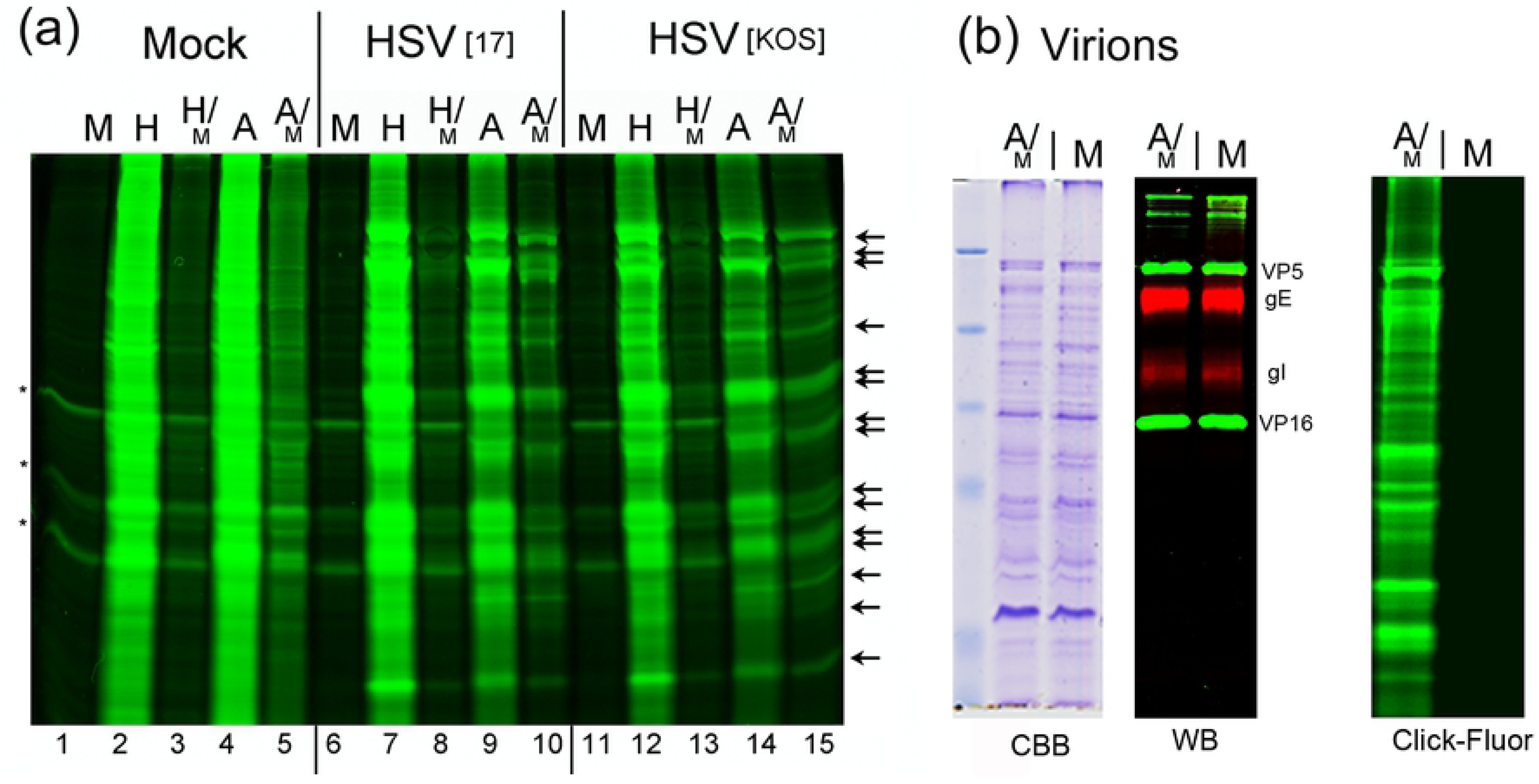
Analysis of bio-orthogonal labelling and comparison of protein abundances in HSV^AHA^ and HSV^wt^ viruses. (a) Cells were mock-infected or infected with HSV-1[17] or HSV-1[KOS] and incubated with combinations of amino acids as indicated in the text from 2-24 hpi. Cells were then lysed and proteins analysed by in-gel fluorescence. Labelled protein profiles are shown for mock-infected (lanes 1-5), HSV-1[17] infected (lanes 6-10) and HSV-1[KOS} infected cells (lanes 11-15). Arrows indicate virus-induced bands. (b) HSV^AHA^ or HSV^wt^ viruses were purified from cells labelled under the combined AHA/Met regimen (A/M) or with normal Met alone (M). Equal aliquots of infectious virus (approximately 10^8^ pfu) were lysed and analysed by SDS-PAGE and total protein staining (CBB) or Western blotting (WB) using representative virion components, VP5, VP16, gE and gI. In parallel equivalent amounts of the viruses were lysed and subject to CuAAC ligation to YnTB, and analysed in in-gel fluorescence (Click-fluor).

To examine this, we produced extracellular HSV virions, after infection with HSV-1 [KOS] using the A/M labelling regime. We compared such preparations (HSV^AHA^) with those produced in standard methionine only medium (termed HSV^wt^). Virions were purified by centrifugation through a Ficoll cushion. Equal infectious pfu were then analysed either by: SDS-PAGE and total protein staining (Fig 2b, CBB); Western blotting of representative structural proteins VP5, VP16 and glycoproteins gE and gD (Fig 2b, WB); or by CuAAC ligation and in-gel fluorescence (Fig 2b, Click-Fluor). Both the total protein and Western blot profiles show virtually identical amounts of the various structural proteins in each virus preparation (lanes A/M and M for HSV^AHA^ and HSV^wt^ respectively). Considering the preparations were standardised on the basis of infectivity, and have equivalent amounts of major structural proteins, the results also indicate that the overall particle/pfu ratios of HSV^AHA^ and HSV^wt^ were similar. Critically, when equal amounts of virion lysates were examined after CuAAC ligation to YnTB, a broad profile of fluorescent proteins was observed for HSV^AHA^ in contrast to the complete absence of signal for HSV^wt^. These data demonstrate the high specificity of CuAAC ligation for proteins incorporating AHA and confirm that AHA was incorporated into a broad spectrum of structural proteins in HSV^AHA^, enabling detection by in-gel fluorescence.

### Mass spectrometry analysis of HSV^AHA^ virions

We next scaled up virus production in bulk roller flasks and analysed virion composition by mass spectrometry. HSV^AHA^ and HSV^wt^ virions were then purified and titrated, again with similar yields (Fig 3a). Again similar levels of representative capsid and envelope-associated HSV proteins were detected in equivalent infectious units (Fig 3b). HSV^AHA^ and HSV^wt^ particles were then lysed, trypsin digested and proteins quantified by shotgun proteomics. Virion components, such as VP5 (MCP), VP1-2, VP16, gB, gD together with other structural proteins were reliably detected and quantified in both preparations. The results are illustrated in Fig 3c, and further tabulated in Table S1 (Preparation 1) comparing protein abundances between HSV^AHA^ and HSV^wt^ and demonstrating very good co-linearity across protein species (Pearson correlations > 0.96 and R^2^ values > 0.93). MS analysis of independent preparations of HSV^AHA^ and HSV^wt^ (S1 Table Preparations 2,3 and Fig S1), again demonstrate very good co-linearity. Altogether the data above provide convincing support that the AHA-labelling regime had no significant effect on either the yield, infectivity or the overall protein composition of HSV.

**Fig 3.**
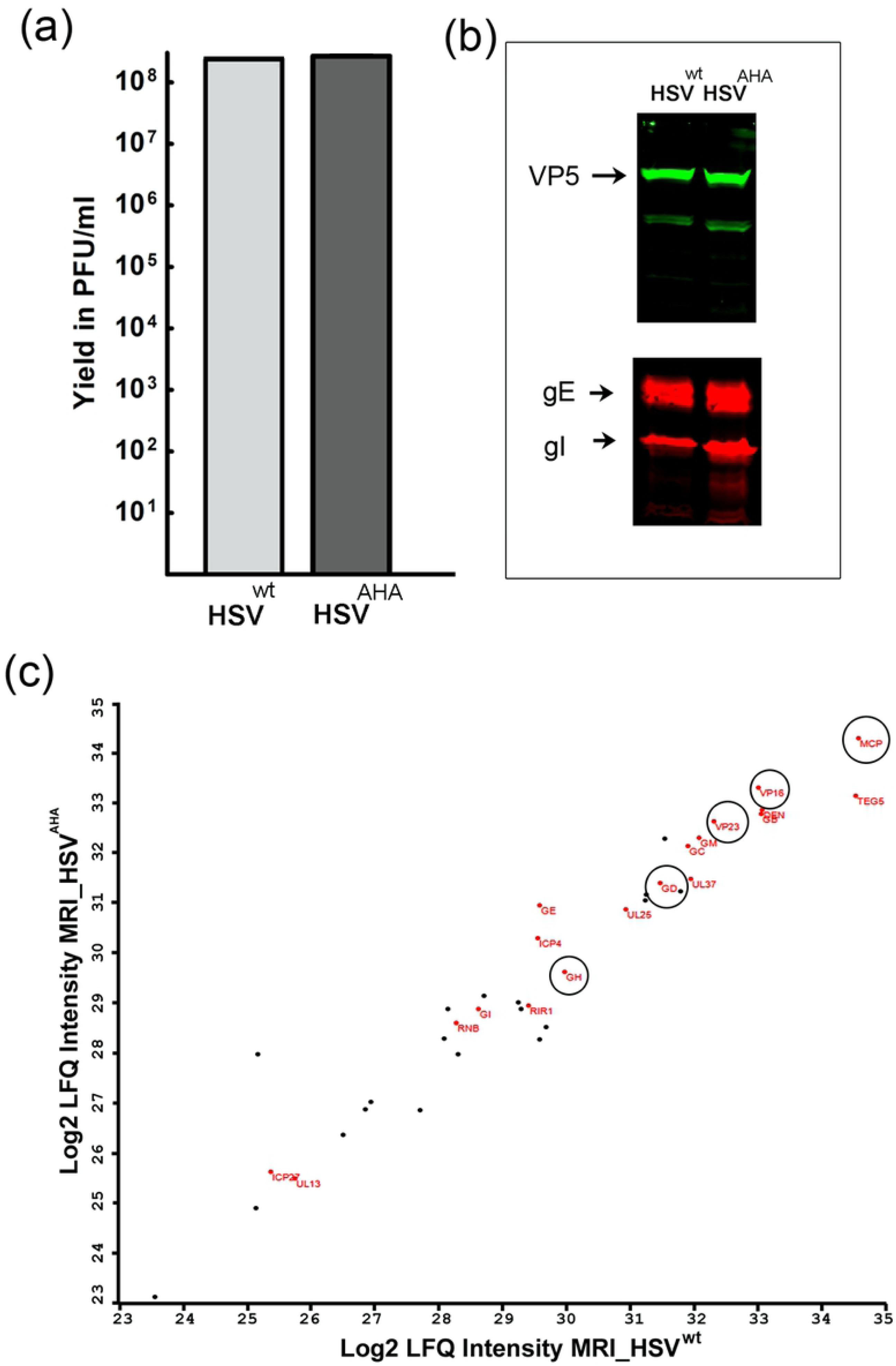
Quantitative comparison of protein abundances in HSV^AHA^ and HSV^wt^ viruses. (a) Large scale virus stocks were made using multi-round replication and the A/M regime or Met alone. Infections were performed in triplicate for each condition. Yields were quantified by plaque titration and plotted as the mean of the triplicates. (b) Equivalent samples of purified extracellular viruses were analysed by Western blot using antibodies to VP5, gE and gI. (c) Equivalent samples normalised based on infectivity were lysed, digested with trypsin and analysed by LC-MS/MS. The results are plotted as the correlation between Log_2_ LFQ Intensities for proteins from HSV^AHA^ (Y-axis) and HSV^wt^ (x-axis). Proteins where Met→AHA substitution sites were detected are highlighted in red. Data for Log_2_ LFQ Intensities are in Table S1.

The in-gel fluorescence shows that a broad range of virus proteins incorporate AHA but does not reveal their specific identities. To examine this we analysed the tryptic peptides generated from HSV^AHA^, unequivocally identifying 77 Met→AHA substitution sites on 22 different viral proteins (Table S2, also indicated in red lettering in the analysis of total abundance, Fig 3c). Most AHA-modified peptides in HSV^AHA^ were detected in abundant or larger proteins for which a higher overall number of peptides were identified. We also wished to estimate the degree of AHA incorporation into HSV proteins. In principle this relates, for any methionine-containing peptide, to the fraction that the Met→AHA form represents of the total amount of that peptide. In practice for reasons outlined in the supplementary information, this question is very difficult to answer. However we attempted to address this indirectly, using AHA labelling with simultaneous pulse-SILAC based quantitation, to supply a surrogate measure of relative amino acid incorporation. The protocol is described in detail in methods and S2a Fig and the results illustrated S2b Fig. Each vertical bar on the X-axis represents an individual HSV virion protein and the Y-axis represents a measure of % AHA incorporation for that protein. In S2c Fig, the % incorporation was binned into 10% brackets and the number of HSV proteins for each bin plotted. Using this surrogate measure, HSV proteins present in extracellular virions incorporated AHA to an extent ranging from approximately 23% to 88% (see also S3 Table).

### Visualisation of individual HSV^AHA^ particles and by CuAAC ligation

We next addressed whether AHA incorporation would allow direct visualisation of individual virus particles by CuAAC ligation of a fluorophore to structural proteins within the particle. Samples of HSV^AHA^ or HSV^wt^ were applied onto borosilicate coverslips, fixed and processed for simultaneous particle detection by immunofluorescence, using a monoclonal antibody to the major capsid protein VP5 or, for AHA-incorporating particles, by CuAAC ligation to alkyne-488. VP5 represents a generally accepted marker used to identify HSV capsid containing particles. Typical results are shown in Fig 4a for VP5 (red channel), AHA (green channel) and the merged image. For HSV^AHA^, almost every particle detected by anti-capsid antibody could be detected by alkyne-488 ligation with examples indicated by vertical arrows (Fig 4a, HSV^AHA^). This was completely specific with essentially no capsid of HSV^wt^ being detected by ligation in parallel assays (Fig 4a, HSV^wt^). While all HSV^AHA^ capsid+ve particles were detected by alkyne-488 ligation, the same was not true in reverse. In other words, there was an excess of total AHA+ve particles over capsid containing particles (Fig 4a, indicated by >), more clearly seen in the inset (Fig 4b). All VP5+ve capsids (red channel, circled) were also detected by CuAAC ligation (green channel), while there were also clearly green particles which were not detected by anti-VP5 immunofluorescence, as summarised in quantitative evaluation of the individual particles (Fig 4c).

**Fig 4.**
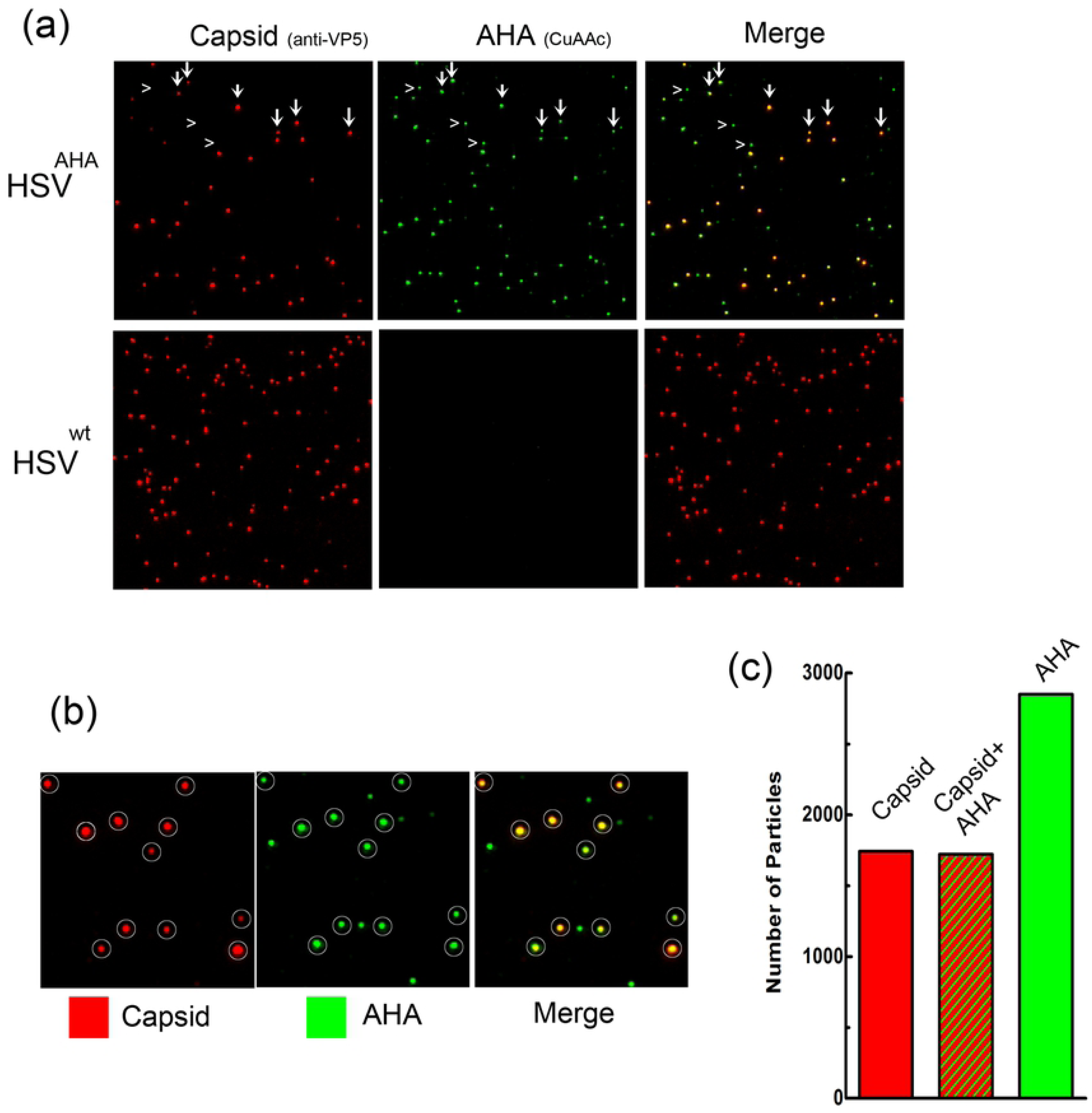
Visualisation of individual HSV^AHA^ particles and by CuAAC ligation. (a) Samples of HSV-1^AHA^ or HSV-1^wt^ were adsorbed onto coverslips prior to processing by CuAAC ligation to Fluor 488-alkyne and immunofluorescence for VP5. Image acquisition was as described in methods. Individual channels are shown for capsid immunofluorescence (red channel) and for CuAAC ligation (green channel), together with the merged image. Vertical arrows indicate representative capsid+ve particles with the same arrows indicated in the AHA and merged channels. Horizontal arrowheads indicate AHA+ve particles which are negative for capsid staining. (b) Enlarged inserts from panel A with capsid+ve particles circled. (c) Quantitative summary of imaging analysis wherein approximately 2800 were identified and scored as capsid+ve (red), AHA+ve (green) or both.

More extensive particle analysis by immunofluorescence versus CuAAC ligation was carried out (Fig 5). In particular it was important for subsequent imaging analysis of HSV^AHA^ entry (see below) that the relative efficiency of individual particle detection was quantitatively estimated. Approximately 1000 HSV^AHA^ individual particles were binned into intensity ranges for each channel (or the zero bin when below the threshold). Of the total, approximately 440 (Fig 5a, population A) scored positive for capsid identification while 615 particles (population B, identified by being positive for the AHA signal), were not significantly positive for VP5 (Fig 5a, population B). The capsid+ve population was sorted on the basis of individual particle intensity (Fig 5b). The analysis indicated a single normally distributed population with the variance in intensity similar to that seen from ours and other laboratories for single particle analysis [26, 27]. Virtually all (>99%) of the capsid+ve particles scored positive for significant AHA signal above threshold. There was no correlation between the signal intensity by anti-VP5 immunofluorescence versus CuAAC ligation, which detects the sum of all available AHA-containing proteins (Fig 5b). The AHA signal of the capsid+ve particles, i.e. population A, also showed a good fit to a normal distribution (Fig 5c). The VP5-ve population tended to show somewhat lower intensities for the AHA signal compared to the capsid+ve/AHA+ve particles (Fig 5d). This likely reflects the observation that capsid proteins are among the main AHA-containing proteins detected by MS (Fig 3 and S2 Table). Over the course of this work, AHA+ve but VP5-ve particles ranged from approximately 1.5 to 3-fold more numerous than the VP5+ve particles. Although it is possible that AHA detection is even more sensitive than immunofluorescence using VP5, and that the excess particles are full virions, we believe this to be unlikely and not in keeping with the data. Rather we propose that these represent the well-recognised HSV L-particles, which contain structural tegument and membrane proteins but lack capsids [28, 29]. The precise characterisation of AHA+ve/VP5-ve particles, which could also include other types of vesicles, is beyond the scope of this current work but is speculated upon in the discussion. With regard to the immediate question, our results convincingly demonstrate that individual VP5+ve capsid containing virions were readily and efficiently detectable by CuAAC ligation and in a highly specific and quantitative manner.

**Fig 5.**
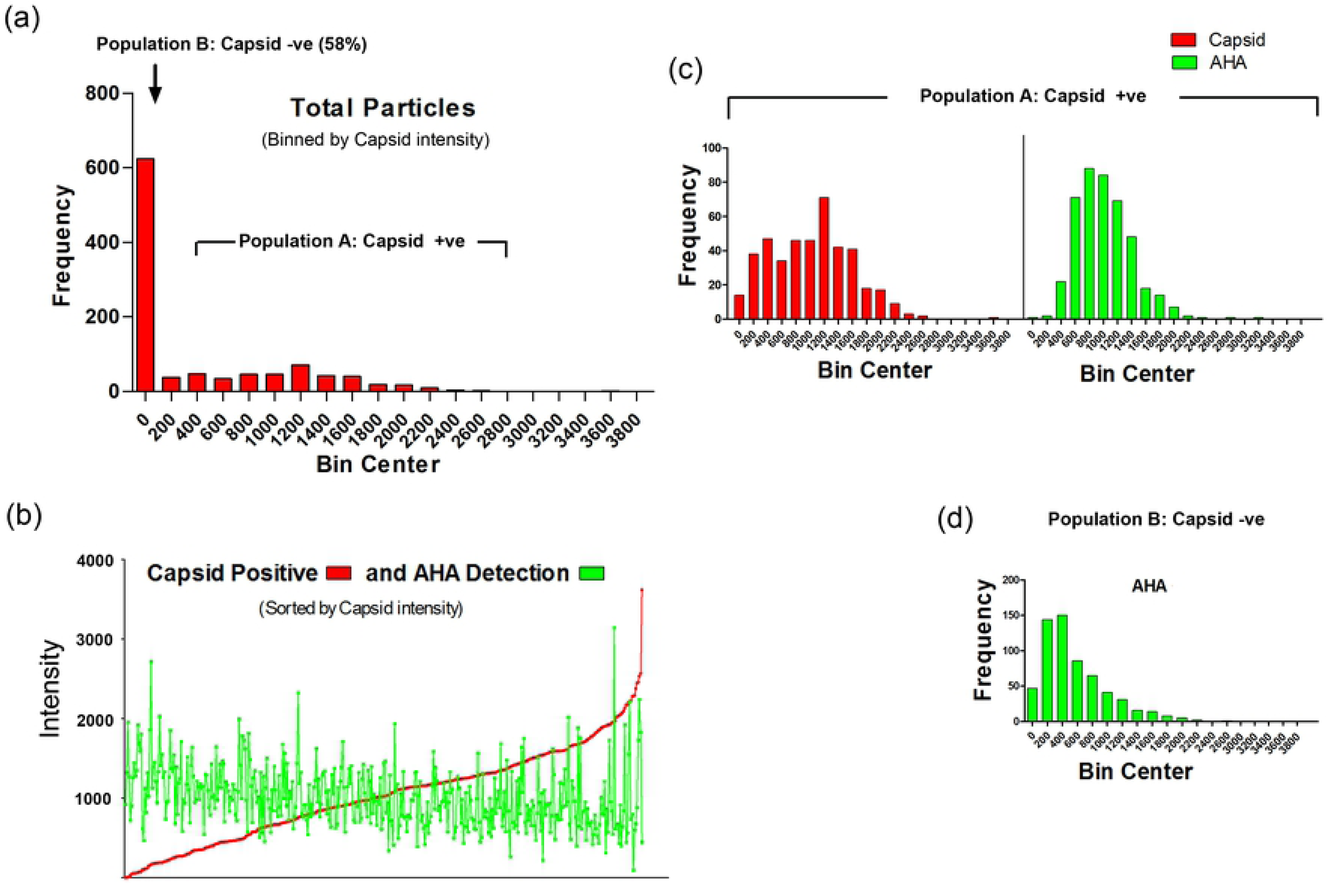
Quantitative analysis of HSV^AHA^ particles by immunofluorescence and CuAAC ligation. Populations of virus particles from images collected as described in Fig 4 were analysed using Image J and a custom plugin [11]. Red (capsid) and green (AHA) intensities were measured for each particle. To be positive, a particle ROI must be at least 1 SD above the background for an equivalently sized ROI for that channel. Frequency distributions of individual particle ROIs were then quantitated and binned. (a) The distribution of all particles scored on the basis of being capsid+ve or -ve. Population B is scored as being capsid-ve, (i.e. below the defined threshold) while all particles in population A are above that threshold and binned into various intensities. (b) Individual population A capsid+ve particles were sorted by increasing capsid signal intensity and the parallel intensity score for each particle for AHA detection plotted. (c) Population A capsid+ve particles were binned into intensity ranges in parallel with the AHA intensity scores across the population. (d) AHA+ve particles that were –ve for capsid detection (population B) were binned into intensity ranges with same axis and bin widths.

### Analysis of early infection by HSV^AHA^ in vivo using CuAAC ligation

We next investigated early virus entry using CuAAC ligation versus conventional immunofluorescence to visualise HSV^AHA^ particles. In addition to the application AHA-labelling in virus detection without antibodies or recombinant fluorescent viruses, we reasoned that analysis of bio-orthogonally modified viruses could reveal processes not readily visualised by conventional means. In particular for example, whether we could discriminate between initially infecting particles, which would be capsid+ve by immunofluorescence and AHA+ve by CuAAC, versus progeny particles which would be detected by immunofluorescence but not by CuAAC.

Cells were infected with HSV^AHA^ at moi 2, where statistically it is expected that a population of cells will initially not be infected [30]. Virus was adsorbed at 4°C (where enveloped viruses bind to cells but cannot fuse) and samples fixed immediately or after shift to 37°C for increasing periods to allow virus fusion and cell entry. Cultures were then processed for immunofluorescence of capsids (anti-VP5, red channel) and by CuAAC ligation to detect AHA+ve particles (green channel). Maximum projection images collected as described in methods.

A typical field at 4°C is shown in Fig 6a and inset (see also examples in S3a Fig). Numerous VP5+ve particles were randomly distributed across the cells. Importantly, every VP5+ve particle was also AHA+ve (individual channels and inset, arrowed particles). There were also AHA+ve particles that were not detected by VP5 immunofluorescence (inset, circled). Quantitative analysis of ca.1500 particles on cell monolayers (S3a Fig) indicates that ∼99% of capsid+ve particles were AHA+ve and that there was an overall excess of approximately 2-fold of AHA+ve/VP5-ve particles over AHA+ve/VP5+ve particles. This data is summarised in Fig S3b,c. By 2 hrs after shift to 37°C, as well as individual particles, clusters could now be observed in many cells (Fig 6b and inset, large arrows). These clusters likely represent the well understood particle congregation and retrograde migration towards the microtubule organising centre [31, 32] or potentially multiple particle uptake within endocytic vesicles [33]. Note that using maximum projections our results do not discriminate between individual particles that have entered cytoplasm and residual particles that may remain bound to the cell surface. Clearly however the cells are infected and initiate virus gene expression with 25-30% positive for VP5, a late product, within 6 hrs of infection (see below). Spatially clustered or coalesced particles reflect cell entry. While individual particles could be resolved within such clusters, the latter were mostly observed as large irregular but spatially restricted regions within the cytoplasm, detected by both VP5 immunofluorescence and CuAAC ligation with broad overlap in the signals (Fig 6b, Fig 7a). However a novel feature of was also observed. We frequently detected large AHA+ve regions that could not be predicted from VP5 localisation and completely lacked anti-VP5 reactivity (example Fig 7b, arrowed). These were never observed at +4°C. In terms of size and location, such regions resembled the AHA+ve/VP5+ve regions (compare Fig 7a and b) but VP5 reactivity could no longer be detected, even though other individual particles in the same cell could be detected as both AHA+ve and VP5+ve (Fig 7b). Furthermore, these large AHA+ve areas could form distended, interlinked cytoplasmic tubules, emanating from an intense clustered centre (example Fig 7c and inset). Within these regions VP5 was present in discrete particles or particle clusters. However the AHA signal was present both within the particles (requiring longer imaging exposure times) but additionally in a diffuse pattern within the tubules that were clearly not particulate in nature and was VP5-ve and ran through the confined cytoplasmic area (Fig 7, panel c inset, arrows positioned for spatial reference points). These extended regions of AHA+ve reactivity could sometimes be correlated with refractile/vesicular regions observed by phase microscopy (Fig 7, panel c, right hand panel). We initially proposed that these extended AHA+ve regions could reflect certain aspects of particle uptake and trafficking and/or coalescence into lysosomal compartments. To examine this latter possibility we performed parallel analysis of the spatial relationship between lysosomes (using the marker LAMP2) and AHA+ve particles or large domains. The results demonstrated that neither individual AHA+ve particles nor AHA+ve macrodomains showed any significant spatial correlation with LAMPs (Fig 8). Nevertheless, previous comprehensive analyses of bio-orthogonal methionine analogues [5–10], and our own results on protein synthesis in HSV infected cells [12, 34], demonstrate that AHA incorporates into proteins during in vivo labelling. Therefore notwithstanding the lack of colocalisation with lysosomal markers, we propose that these AHA+ve macrodomains represent cytoplasmic sites of virion processing or breakdown, releasing proteins, fragments or peptides within membrane-bound compartments with such fragments not being detected by conventional immunofluorescence with antibodies. Possible explanations expanded upon in the discussion.

**Fig 6.**
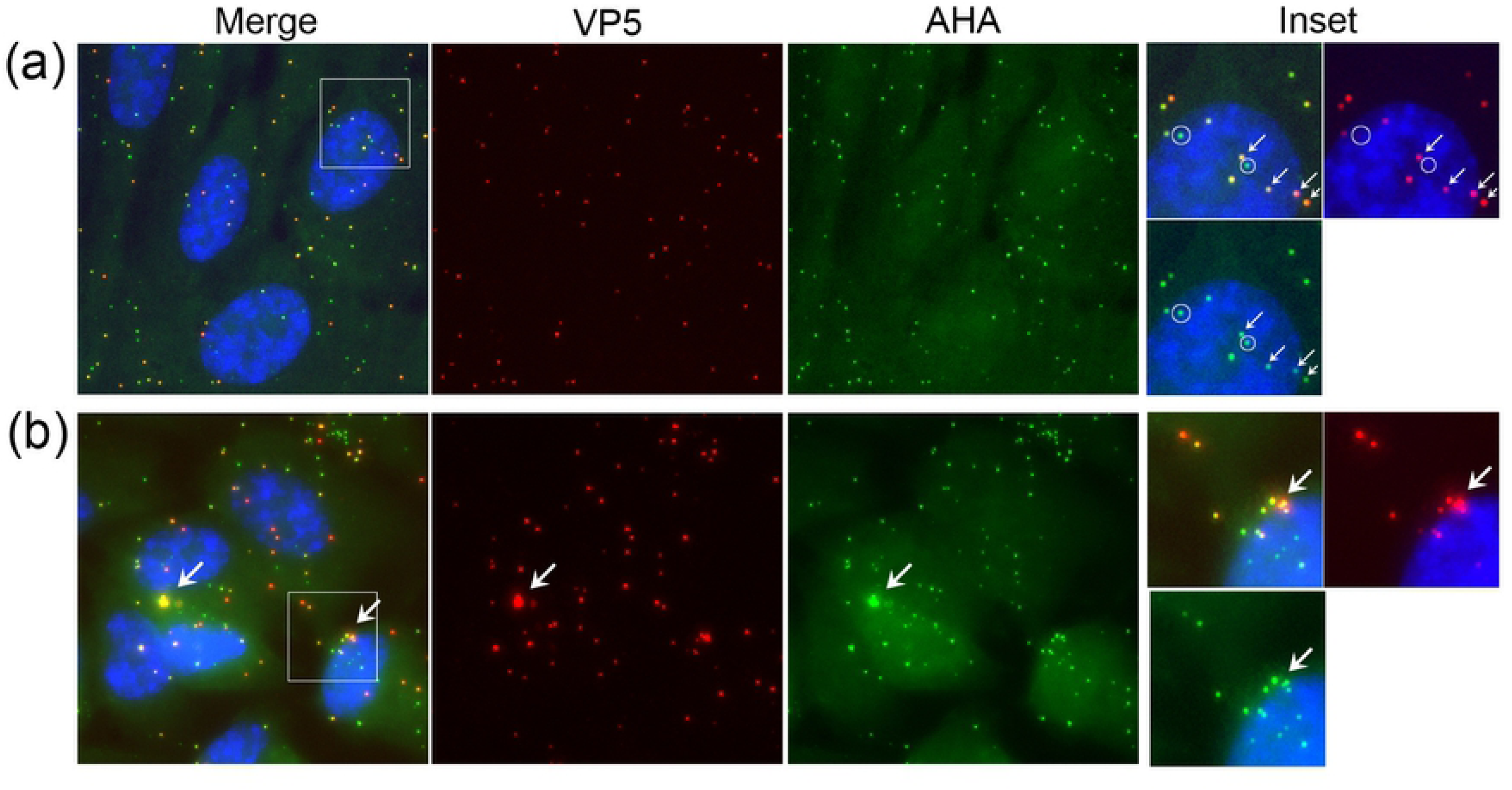
Analysis of HSV^AHA^ cell entry by CuAAC ligation of infecting particles. Cells were infected with HSV^AHA^ and incubated +4°C then fixed immediately (a) or 2 hrs after shift to 37°C to allow virus entry (b). Particles detected by CuAAC ligation versus detection by anti-VP5 capsids immunofluorescence are shown in the green and red channels respectively. The box in the merged image is expanded in the inset right hand side. Arrows and circles are as described in the text.

**Fig 7.**
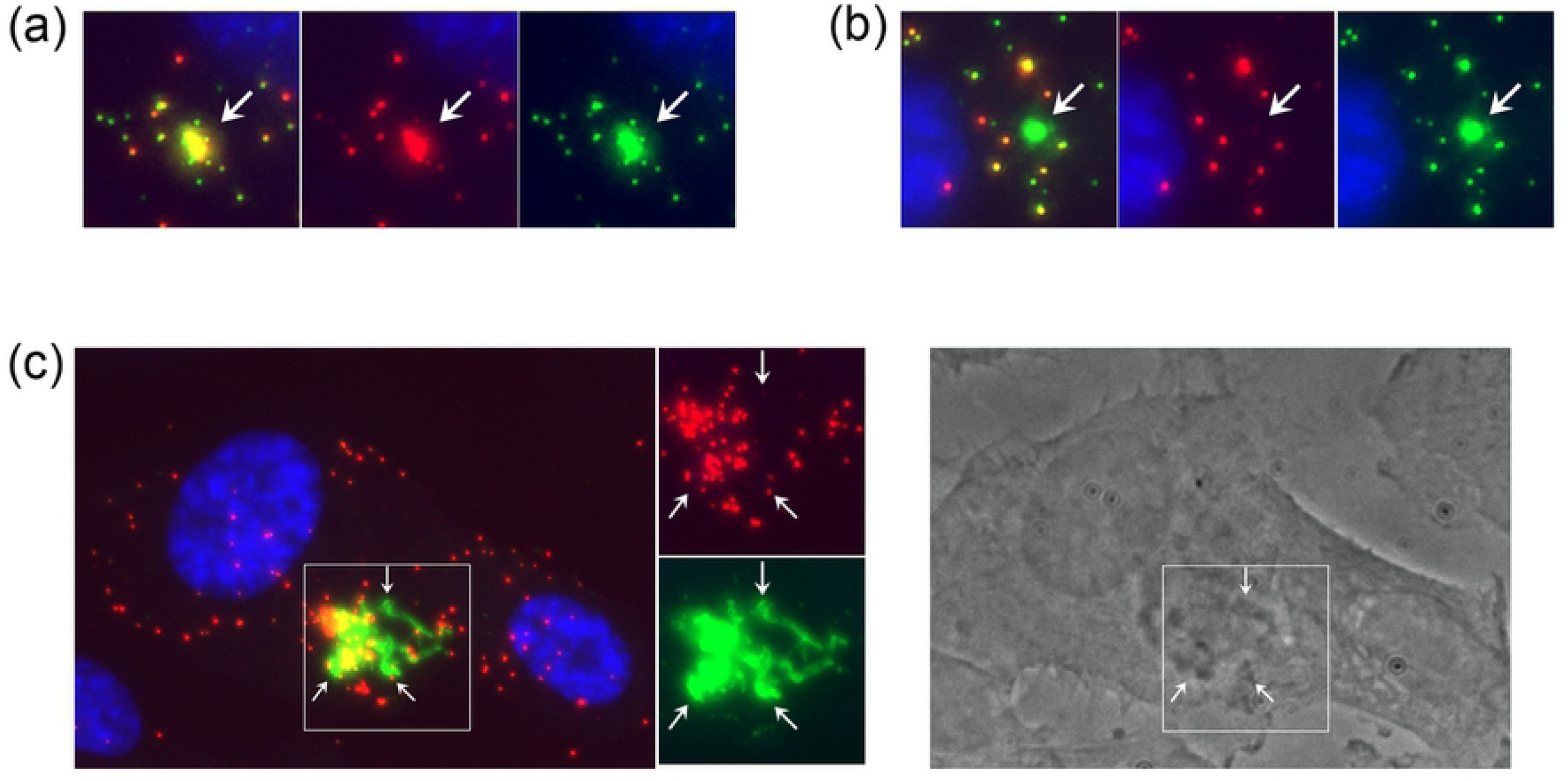
Formation of AHA-containing cytoplasmic macrodomains during entry. Cells were infected with HSV^AHA^ processed at 2 hpi and presented as described in Fig 6. Panels and insets illustrate representative cells showing features discussed in the text. Note in panel c, the high intensity of AHA signal in the central cluster required very low exposure times for spatial resolution, such that the individual VP5+ve particles (red channel) were then poorly detected for AHA. The box in the merged image is expanded in the inset with arrows placed in identical positons for spatial reference in each channel.

**Fig 8.**
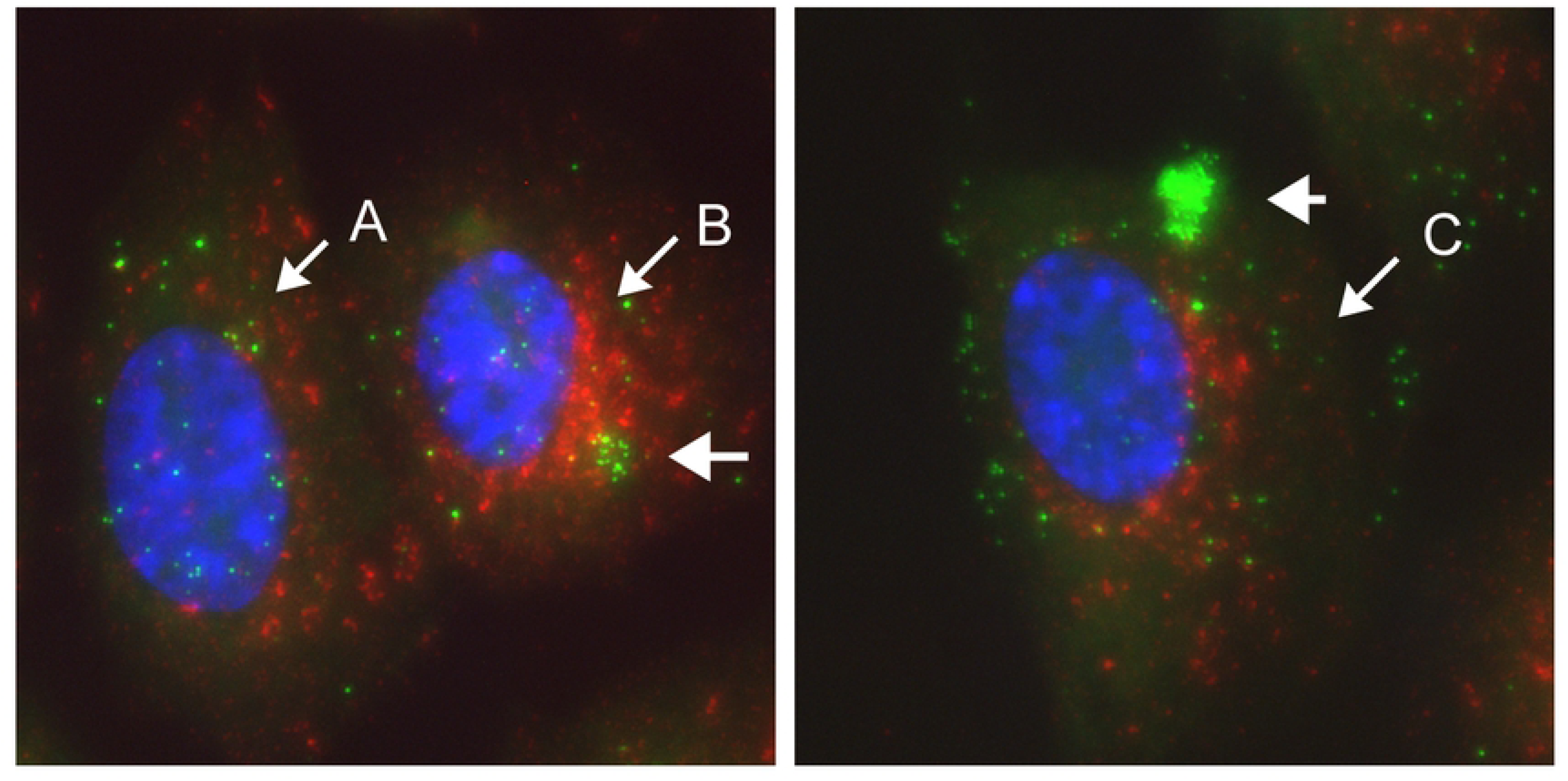
AHA-containing cytoplasmic macrodomains do not colocalise with lysosomal compartments. Cells were infected with HSV^AHA^ processed at 2 hpi by CuAAC ligation for AHA detection (green channel) or immunofluorescence for lysosomal detection using the marker LAMP2 (red channel). Distinct examples of AHA localisation are illustrated Cell A shows a cell with mostly individual particles. In Cell B, the AHA signal forms a tight regionally defined smaller cluster where individual particles could still be resolved and Cell C shows an extremely intense AHA+ve area.

### Discrimination of retrograde, infecting virus from replicated anterograde secondary infections

From approximately 4 hrs, de novo synthesised VP5 was synthesised in an asynchronous manner, distributed within the nucleus and as expected showed no detection by CuAAC ligation while the infecting VP5+ve particles were readily detected (Fig 9, individual channels and inset). By 6 hrs, approximately 20-30% of the cells showed diffuse de novo synthesised VP5 (Fig 10 and S4a Fig). Cells negative for nuclear VP5 (Fig 10a, cell type A) could represent cells which were not yet productively infected or were infected but not yet positive for de novo synthesised VP5. By this time point additional features could now be ascertained. Thus in certain cells where de novo nuclear VP5 was abundant (Fig 10a, cell B), VP5+ve capsids could also be observed in the cytoplasm (inset, small vertical arrows). However these capsids were all AHA+ve and virtually no VP5+ve/AHA-ve cytoplasmic capsids were observed in this class of cell. Note that weaker diffuse/microspeckled VP5 staining in the cytosol represents soluble or subunit protein, readily discriminated from VP5+ve bright defined capsid particles. Thus cell type B represents cells with abundant de novo nuclear VP5, but where cytoplasmic VP5+ve particles do not represent early progeny capsids in the cytoplasm but rather initial infecting virions.

**Fig 9.**
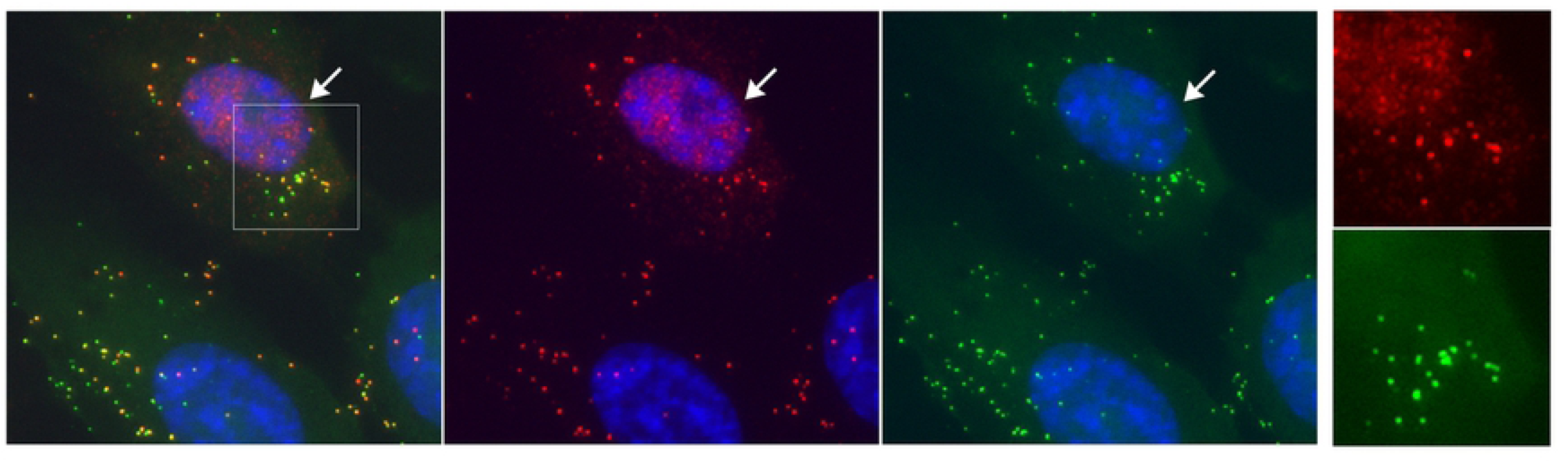
Analysis of HSV^AHA^ cell entry. Cells were infected as for Fig 6 showing cells infected with HSV^AHA^, and fixed 4 hrs after shift to 37°C and processed for VP5 by immunofluorescence or AHA by CuAAC ligation. The box in the merged image is expanded in the inset with the arrow indicating de novo synthesised VP5 in the nucleus.

**Fig 10.**
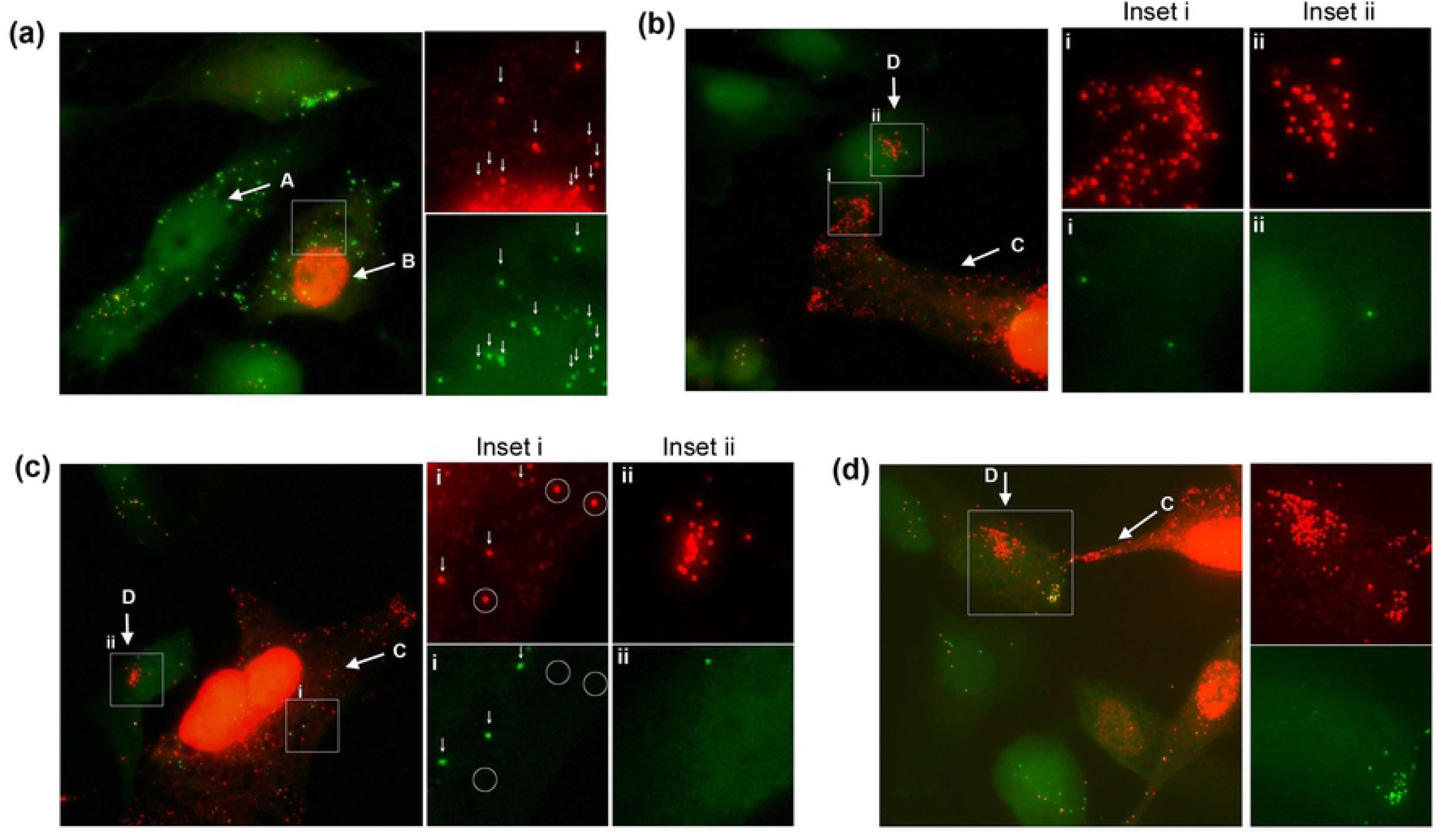
HSV^AHA^ selective identification of infecting versus progeny capsid. 6 hrs after shift (b-e). Particles detected by CuAAC ligation versus detection by anti-VP5 capsids immunofluorescence are shown in the green and red channels respectively. Panels and insets illustrate representative cells showing features discussed in the text. The nuclear signal (blue channel) has been omitted where needed for clarity.

In other cells representing more advanced infection, abundant nuclear VP5 was also observed now together with numerous cytoplasmic VP5+ve particles where the vast majority were now undetectable by CuAAC ligation (Fig 10b, cell C, inset i). Although it could be formally proposed that such particles represented infecting virions that had somehow completely lost their AHA signal, considering their numbers and the fact that VP5 is one of the main AHA-containing proteins, the simplest interpretation is that AHA-ve/VP5+ve capsids represent progeny capsids. Indeed progeny capsids could be discriminated from infecting particles in the same cells. An example (Fig 10c, cell type C, inset i), shows the low frequency VP5+/AHA+ve input particles (vertical arrows) and the VP5+ve/AHA-ve particles (circled) that were much more numerous throughout the remainder of the cytoplasm. Using just the VP5 signal, i.e, by conventional immunofluorescence, no such discrimination could be made. To confirm that VP5+ve/AHA-ve particles were progeny capsids, we examined infection in the presence of phosphonoacetic acid (PAA), an inhibitor which blocks HSV DNA replication and prevents full capsid assembly and normal egress to the cytoplasm. In this case, few cytoplasmic capsids were observed and those present were invariably AHA+ve (S4b Fig and inset). The results strongly support the conclusion that VP5+ve/AHA-ve particles seen in the normal course of HSV^AHA^ infection represent progeny capsids and that progeny and initial infecting capsids can be effectively discriminated.

Our results also allowed identification of an additional cell type (Fig 10b-d; cell type D). In these cells, numerous VP5+ve particles could be now be observed exclusively in the cytoplasm. However type D cells had no nuclear VP5 signal that would indicate de novo synthesised VP5. Moreover in type D cells, the numerous VP5+ve capsids frequently congregated in tightly clustered areas, just adjacent to the nucleus. Importantly, in type D cells these capsids were also invariably AHA-ve (Fig 10b-d; insets ii). Type D cells were never observed in the presence of PAA. Altogether these considerations demonstrate that the capsids in type D cells, rather than representing slower initiating infections, represent secondary infected cells, infected with progeny viruses assembled in primary infected cells. Furthermore we could also observe that type D cells could be dually infected in a localised and spatially distinct manner regarding primary versus secondary infections (Fig 10d). Initial infecting VP5+ve/AHA+ve particles were observed clustered in a peripheral region of the cell (Fig 10d, inset, red+green particles, located lower right) while VP5+/AHA-ve particles, (infecting from the adjacent cell C), were present in high numbers in a tightly localised manner and in a distinct perinuclear region of the same cell (cell D and inset, red only particles, located upper left). These data indicate that virus had already replicated assembled and infected adjacent cells by 6 hrs and could be selectively spatially organised in the cell distinct from the initial infection. The results reveal aspects of the spatiotemporal landscape of infection that would be difficult to demonstrate with conventional methods as discussed further below.

## Discussion

We demonstrate the replication and assembly of a complex enveloped DNA virus, HSV, in the presence of non-canonical bio-orthogonal methionine analogues. HSV encodes over 70 proteins including very abundant structural proteins that are recruited in 100s to 1000s of copy numbers per virion [27, 30, 35]. An optimized AHA-labelling regime allowed production of normal yields of HSV^AHA^ particles that were fully infectious and characterised by unaltered protein composition compared to unmodified HSV^wt^ particles, as measured both by WB analysis and quantitative proteomics. Furthermore, mass spectrometry revealed Met→AHA substitution sites in 30% of all HSV proteins (37% of HSV proteins identified in this study), mostly in abundant or larger proteins. Using a surrogate pulse SILAC approach, we estimate that proteins detected in extracellular virions could have AHA incorporation rates ranging up to 50%, indicating that rather than small percentages or fractions of a percent, AHA was relatively efficiently incorporated into HSV structural proteins. We note also that lack of detection of Met→AHA substitutions does not necessarily mean lack of AHA incorporation and could simply reflect low abundance or disadvantageous physicochemical properties of tryptic peptides (e.g. extreme hydrophilicity or hydrophobicity, low solubility, sub-optimal size). We also identify sites of Met→AHA substitution for certain of AHA-containing proteins. Thus, notwithstanding changes in the translational apparatus and the demands on synthetic capacity of high copy number structural proteins, bio-orthogonal amino acids are extremely well tolerated and incorporated during virus protein synthesis and assembly of infectious virions. Previous work has also indicated that bio-orthogonal amino acids can be incorporated into the non-enveloped Adenovirus 5 capsids without compromising infectivity with several capsid proteins identified by mass spectrometry, although no comparative proteomics or entry studies were carried out [19].

At an individual particle level, we demonstrate that virtually every VP5 containing HSV virion was also detected by CuAAC ligation. Two specific features of the analysis of virion cell entry of HSV^AHA^ using CuAAC ligation and imaging merit discussion. As well as individual VP5+ve/AHA+ve particles, we observed large intracellular areas of dense aggregated regions, sometimes with individual particles resolved within them. These aggregated areas were both VP5+ve and AHA+ve, but the two signals could also segregate differentially within the same overall spatial area of the cell. Thus we frequently observed extremely intense AHA+ve areas that had little and sometimes no VP5+ve signal discernible within them. Such regions were also observed as intense extended interlinked tubules enveloping and surrounding areas of dense VP5+ve coalescing particles.

We interpret these results as follows. Many infecting particles may remain as individual resolved particles in the cytoplasm or coalesce in a concentrated manner, e.g. around the microtubule organising centre. However in many cells, possibly though not necessarily reflecting particle number, particles may also be taken up by an endocytic or macropinocytic route, particularly in keratinocytes, which are known to support endocytic virus entry [33, 36]. We propose that the large coalesced AHA+ve areas represent defined and membrane-restricted regions reflecting such uptake with subsequent intracompartmental processing. Initially these regions would be both VP5+ve and AHA+ve, accepting there will also be an excess of VP5-ve/AHA+ve particles. Although warranting future investigation, we no evidence for any spatial segregation of VP5+ve/AHA+ve particles from VP5-ve/AHA+ve particles during entry.

We propose that within these areas, virions and particles are disassembled or partially degraded into proteolytic fragments and peptides. The massive increase in AHA signal is because, while we clearly show that proteins are accessible to CuAAC ligation within intact particles, particle disruption or proteolysis, within a membrane confined area, would expose numerous AHA sites not available in the intact particle, yielding a dramatic increase in fluorescence intensity while at the same time diminished or even complete loss of detection of specific epitopes. While these larger intense areas of virion processing were seen in a subpopulation of cells, they could represent an important aspect of cell-cell heterogeneity during infection with the intense AHA+ve regions being involved in aspects of the general cell response to infection, antigen presentation or paracrine signalling. Our results convincingly show that such regions did not colocalise with lysosomes. Future work will explore whether they co-localise with other endogenous markers or represent completely novel virus induced compartments and any correlation with individual cell responses to infection.

Another aspect of particle detection by CuAAC ligation was the presence of AHA+ve particles that were not detected by VP5 immunofluorescence. Consistent with our general understanding of HSV infection, we propose that such particles represent L-particles which assemble abundant tegument proteins and are enveloped but do not contain capsids. However, AHA+ve particles could also include other particle types, including exosomes, exported from infected cells [37]. To examine the origin of AHA+ve/VP5-ve particles it will be necessary to carry out biochemical or affinity purification, although separation of L-particles from true exosomes has not been reported and may prove difficult. Nevertheless, AHA incorporation may offer new opportunities to examine extracellular vesicle biology because of current compounding features in exosome analysis. These include their presence in calf serum used in virtually all cell culture experiments and the fact that most cells are thought to continually export and may reimport exosomes, rendering it difficult to discriminate origins, directionality and variations in exosome assembly, release and uptake. AHA pulse-labelled extracellular vesicles would allow refined spatial analysis because while all exosomes might be positive for a defining marker, only exosomes from an originating, pulse-labelled cell, would be AHA+ve by CuAAC ligation. This would facilitate spatial discrimination and analysis of the kinetics of transport and changes in protein constituents.

A final aspect from analysis of HSV^AHA^ is the ability to discriminate between infecting virions and progeny virions. For successful infection HSV moves in a retrograde manner towards the cell centre and nuclear membrane. With HSV^AHA^ such virions are VP5+ve/AHA+ve while progeny particles are VP5+ve/AHA-ve. This discrimination virtually absolute, because of the very high selectivity of the CuAAC reaction with no significant signal detected for control HSV^w/t^ particles. When there are comparatively high numbers of capsids in the cytosol it is possible, by immunofluorescence alone, to say that the majority of these are likely to be progeny capsids but it is impossible to discriminate individual progeny from initially infecting particles by immunofluorescence alone. This can be important e.g. in the examination of retrograde and anterograde transport within neuronal axons where it can be difficult to discriminate infecting particles and progeny particles. Moreover, using the AHA status of VP5+ve capsids we could identify cells, even within six hours of infection that clearly represented second round infection by progeny virus, frequently in a spatially restricted perinuclear area. We observed that cells could be dually infected with numerous AHA+ve/VP5+ve infecting capsids present together with AHA-ve/VP5+ve capsids, in distinct spatially restricted areas of the same cell. Although it is generally accepted that second round infections are typically high multiplicity, our results indicate that second round infections could take place extremely rapidly and could involve additional processes. The spatial segregation of first and second round virus in the same cell could for example involve increased efficiency of vectoral transport of second round progeny capsids by manipulation of processes in naive cells surrounding infected cells, or by spatially restricted assembly of particles in the primary infected cell. Such questions together with the use of bio-orthogonally modified virions to investigate the biochemical composition or relative protein exposure during infection will be the subject of future investigations.

Altogether our results help demonstrate both the complexity in early entry processes and the potential for bio-orthogonally modified viruses for future investigation of the transport and fate of infecting particles. Bio-orthogonally modified viruses are not limited to those containing noncanonical amino acids and previous results have also demonstrated the ability to assemble viruses with bio-orthogonally sugars and the possibility of altering host range [38, 39]. Together with this work reporting a tractable pipeline for production of a complex, enveloped virus with bio-orthogonally modified proteins and previous work on HSV with bio-orthogonally labelled genomes, the opportunities for these techniques in different viruses and different fields of infection biology are considerable.

## Materials and Methods

### Chemicals and reagents for BONCAT and CuAAC

L-azidohomoalanine (AHA) was from Iris Biotech GMBH; L-homopropargylglycine (HPG) was from Chiralix BV; CuSO4, sodium ascorbate (NaAsc), EDTA, Alexa 488-alkyne, Cy3-alkyne, tris(2-carboxyethyl)phosphine (TCEP), tris(hydroxypropyltriazolylmethyl)amine (THPTA), tris(benzyltriazoylmethyl)amine (TBTA), and aminoguanidine (AG) were all from Sigma-Aldrich. Azido-TAMRA-Biotin (AzTB) and Alkynyl-TAMRA-Biotin (YnTB) were synthesized as described previously [40].

### Cells, virus and infection

RPE-1, a human telomerase immortalised retinal pigment epithelial line was passaged in Dulbecco’s modified minimal essential medium (DMEM)-F12 (Sigma) supplemented with 200 mM glutamine, 10% fetal calf serum (FCS) and penicillin/streptomycin. Virus strains were laboratory standards HSV-1[17] and HSV-1[KOS]. Routine plaque assays were performed in duplicate in the presence of 1% pooled neutralising human serum (Sigma). Single-step infections were performed at moi 5 or 10 pfu/cell and multi-round infections at a moi of 0.005 pfu/cell. For routine BONCAT labelling of proteins in cell culture, DMEM without Met, cysteine (Cys) and glutamine (Gln) (Sigma) was used, supplemented as necessary. For production of BONCAT-labelled HSV, we used custom medium (Dundee Cell Products) comprising DMEM-F12 lacking Met, leucine (Leu), arginine (Arg), and lysine (Lys) supplemented as described in the text. In dual AHA pulse-SILAC experiments, ^13^C6,^15^N4-Arg and ^13^C6,^15^N2-Lys (CK Isotopes) and SILAC-certified dialysed FBS (ThermoFisher Scientific) were used as supplements.

### Cell proliferation and viability

RPE-1 cells (1.5×10^5^) were plated in triplicate and grown in Met-depleted media supplemented with one of the following: AHA, HPG, Met or with mixtures of the noncanonical amino acids and Met at concentrations indicated. After 24 hrs the cells were trypsinised and evaluated for cell numbers and viability by trypan blue exclusion, enumerated using a Countess Automated Cell Counter (Invitrogen).

### Optimisation of production of HSV labelled with noncanonical amino acids

RPE-1 cells were infected with HSV-1[17] at a moi of 5 in serum-free media. Media were removed and replaced by media supplemented with 2% serum. Approximately 0.5 hr prior to the beginning of the labelling period, the medium was replaced with Met-free media supplemented with 2% dialysed serum to deplete the cells of natural methionine. Met-free media, supplemented with 2% dialysed serum and the indicated concentration of Met, AHA, HPG, or various mixtures thereof, were then added and cells incubated for the times indicated. The incubation medium was removed and infectivity of extracellular viruses determined by standard titration methods. To examine active protein synthesis in parallel, where indicated, infected cells were washed with PBS, lysed in 2% SDS in PBS, and subjected in solution to CuAAC ligation to YnTB or AzTB. Lysed samples were then analysed by SDS-PAGE and in-gel fluorescence imaging (see below, Click chemistry and in-gel fluorescence imaging).

To compare AHA-labelling efficiency and infectivity of viruses, RPE-1 cells were infected with HSV-1 at either moi 5 or at moi 0.005. For single-step infections, labelling medium, consisting of Met-depleted medium supplemented with AHA (1 mM, 2% dialysed serum), was introduced from 9 until 24 hpi. Prior to that, at 8.5 hpi cells were incubated for 30 min with Met-depleted medium supplemented with 2% dialysed serum. For multiple-round infections, we optimised conditions as described in the text. Labelling medium, consisting of Met-depleted medium supplemented with Met (0.02 mM) or Met (0.02 mM) plus AHA (0.6 or 1.2 mM) and 2% of dialysed serum, was introduced from 2 until 72 hpi. At the end of the labelling period, extracellular virus particles were clarified from cellular debris by low speed centrifugation (1,000 ×g, 10 min). Aliquots were then assayed for infectivity of extracellular viruses by standard titration methods. In parallel, aliquots were concentrated by high speed centrifugation (16,600 ×g, 1.5 hrs) to pellet the virus, lysed in 2% SDS in PBS (100 μl) and sonicated briefly (2 × 5s) in an ultrasonic bath. The viral lysates were then subjected to CuAAC ligation to YnTB, followed by SDS-PAGE and in-gel fluorescence analysis.

For large scale preparations HSV^AHA^ and HSV^wt^ were produced in parallel by infecting RPE-1 cells in roller bottles (approximately 2 × 10^8^ cells) with HSV at moi 5 (single-step infections) or moi 0.005 (multi-round infections). After labelling and virus production, the virus containing supernatant was used to purify particles and the cell pellet retained frozen at -80°C. Supernatant virus was pelleted in a Sorvall RC-5C Plus Superspeed Centrifuge with a SS-34 rotor (19,000 rpm, 1.5 hrs). The virus pellet was resuspended in 4 ml PBS and virus subsequently pelleted through a 35% sucrose cushion using a Sorvall WX Ultra 80 centrifuge with an SW55-T1 rotor (23,000 rpm, 1 hr). The supernatant was discarded and the pellet was resuspended in of PBS (500 μl per each roller bottle used for virus production). The resulting suspensions were clarified by a low speed centrifugation (3000 rpm, 5 min) and then by filtration through a Millex-HV 33 mm filter unit (PVDF 0.45 µm). Aliquots of these filtered samples were diluted with PBS to 3 ml and further purified by pelleting through a 2 ml cushion of 10% Ficoll using a Sorvall WX Ultra 80 centrifuge with an SW55-T1 rotor (16,600 rpm, 2 hrs). Total protein profiles of concentrated HSV^AHA^ and HSV^wt^ viruses were visualised after lysis of samples (5×10^8^ pfu) in 2x Laemmli buffer, SDS-PAGE and Coomassie Blue staining. Labelled protein profiles were visualised by lysis of samples, CuAAC ligation to YnTB, SDS-PAGE and subsequent in-gel fluorescence analysis. Relative levels of specific individual proteins in HSV^AHA^ and HSV^wt^ samples were also investigated by western blotting and by mass spectrometry (see below). For western blotting primary antibodies were as follows; monoclonal mouse anti-VP5 capsid protein(1:2500, Virusys) and rabbit anti-gE/I anti-sgE/I envelop protein (1:1000, kindly supplied by Dr Todd Wisner) with secondary antibodies (when imaged with LI-COR Odyssey imager (LI-COR), goat anti-mouse IgG Dylight 800 conjugate (1:10000, ThermoFisher Scientific) and goat anti-rabbit IgG Dylight 680 conjugate (1:10000, ThermoFisher Scientific).

### AHA-labelling for standard analysis of protein synthesis in infected cells

RPE-1 or Vero cells were infected with HSV-1[17] at moi 5 or mock-infected in serum-free media for 1 hr. The medium was removed and replaced by media supplemented with 2% serum. 30 min prior to the metabolic labelling, incubation medium was removed and replaced by medium lacking methionine containing 2% FBS. This medium was then removed and replaced by Met-free medium supplemented with AHA (1 mM) and 2% dialysed FBS. Where indicated, in certain controls, where cells were incubated with AHA in the absence or presence of 100 μg/ml cycloheximide (CHX), CHX was added to the cells 1 hr before pulse-labelling with AHA. At the end of the pulse, the cells were washed with ice-cold PBS (2×) and either fixed with 4% paraformaldehyde in PBS (for image analysis) or harvested in 2% SDS in PBS (for biochemical analysis). For the latter samples were subjected to click chemistry in solution followed in-gel fluorescence imaging.

### Click chemistry in solution and in-gel fluorescence protein analysis

Samples of cell extracts or virus preparations were diluted with PBS to a final SDS concentration of 1% and sonicated using a Braun Ultrasonicator with microprobe (2 × 2 s). Protein concentrations were determined using a Nanodrop Lite spectrophotometer (Thermo Fisher Scientific) and equal protein amounts (typically 100 μg) subjected to CuAAC ligation to AzTB [40]. Mixtures for ligation were prepared by adding reagents in the following order: AzTB (1 μl, stock solution 10 mM in DMSO, final concentration 0.1 mM), CuSO4 (2 μl, stock solution 50 mM in water, final concentration 1 mM), TCEP (2 μl, stock solution 50 mM in water, final concentration 1 mM), TBTA (1 μl, stock solution 40 mM in DMSO, final concentration 0.4 mM). Following the addition of the click mixture (6 μl/sample), the samples were mixed by vortexing for 1 hr at 25°C, and the reaction was stopped by addition of EDTA (2 μl, stock solution 500 mM in water, final concentration 10 mM). Proteins were precipitated by addition of chloroform and methanol in a ratio of 0.25:1:1 v/v/v chloroform/methanol/sample. Precipitated proteins were pelleted by centrifugation (13,000 × *g*, 10 min), washed twice with methanol (1 ml), and air dried. For in-gel fluorescence analysis of AHA-incorporation into purified viruses, the mixtures were spiked with 20 μg of unlabelled total cell lysate to aid efficient protein precipitation. The pellets were then reconstituted in sample loading buffer, boiled for 5 min and loaded on a 12% SDS-PAGE gel. Following electrophoresis, gels were fixed in 40% MeOH, 10% acetic acid, 50% water, and washed with water. In-gel fluorescence was recorded on a Typhoon FLA 9500 imaging system, typically followed by Coomassie Brilliant Blue staining for total protein visualisation and images recording on an Epson Perfection 4990 scanner.

### Simultaneous detection of purified HSV^AHA^ particles by CuAAC cycloaddition and immunofluorescence

Coverslips were washed in ethanol and then PBS. Virus was diluted in PBS and approximately 6 × 10^5^ pfu (in 5 µl) was bound onto the coverslips for 15 min. Viruses were then fixed in 4% PFA in PBS for 15 min, permeabilised in 0.5% Tx-100 and processed for detection by cycloaddition of the alkyne-fluorphore, (Fluor 488-alkyne). After the reaction, samples were made to 1 mM EDTA to stop any further reaction. Samples were then processed by immunofluorescence using an anti-VP5 monoclonal (1:400, Virusys). Secondary antibody was Alexa-Fluor 546 goat anti-mouse IgG (1:750 Life Technologies). For quantitative analysis maximum projections were captured using the Image Pro Plus Stage-Pro function and Z-stacks were obtained with 10 slices at 0.2 µm intervals. We used Image J and a customised plugin based on the find maxima protocol as previously described [11]. The plugin uses find maxima and places an identical sized ROI centred on the maxima with user configurable diameter to encompass virus particles. Maxima with too close a spatial overlap or at an image edge are excluded by the protocol and can be further excluded manually before quantitation. In practice this had a limited effect given the largely monodisperse nature of analysed particles. Red (capsid) and green (AHA) intensities were measured for each ROI. Mean and standard deviation (SD) background intensities were calculated separately for the red and green channels from the area outside the identified ROIs and normalised for ROI area. Maxima–based ROIs were then compared separately against the mean background for each channel and categorised using a threshold the default of which was the mean channel background plus 1 SD. Thus, to be categorised as a red (capsid) positive particle, that particle ROI must be not only be above the mean background ROI in the red channel but at least 1 SD above that background. Frequency distributions of individual identified particle ROIs were then quantitated, calculating the bin width using the Friedman-Diaconis criteria for interquartile-ranges [41, 42]. The same bin width was used for both channels in the Figure for ease of comparison of the distributions. Gaussian distributions were fitted to each channel frequency data using Image J curve fitter.

### Tracking of HSV^AHA^ infecting particles during early infection by click chemistry of virion proteins

RPE-1 cells were seeded on glass coverslips in 12-well plates (2 × 10^5^ cell/well). Cells were pre-chilled at 4 °C, infected with HSV-1^AHA^ (moi 10) in serum-free medium and incubated for a further 1 hr at 37°C to allow virus entry. Cells were then washed, fixed in 4% PFA in PBS (15 min) and processed for simultaneous immunofluorescence and CuAAC cycloaddition. In this case, fixed cells were permeabilised with 0.5% Triton X-100 for 5 min, washed twice with PBS, then processed by CuAAC in reaction buffers containing CuSO4 (1 mM); THPTA (1 mM), NaAsc (10 mM); AG (10 mM); and Fluor 488-alkyne (10 μM) in PBS pH 7.4. Reaction were performed for 2 hrs at 25°C in the dark. After removal of the buffer, cells were washed with PBS (3x). Samples that were to be further processed by immunofluorescence, were blocked and treated for simultaneous immunofluorescence and image capture as above.

### MS sample preparation, instrumentation, and software

Equal aliquots of purified infectious HSV^AHA^ and HSV^wt^ viruses (2×10^7^ pfu) were lysed in PBS buffer (500 µl) containing 2% SDS and sonicated (2 × 5s). Proteins were precipitated by an addition of methanol (500 μl) and chloroform (125 μl), and vortex-mixing. Protein pellets were isolated by centrifugation (17,000 × *g*, 10 min), washed (3x) with methanol (0.5 ml), and air dried. The pellets were then reconstituted by addition of 100 µl of 50 mM ammonium bicarbonate (AMBIC) and 5 µl of 100 mM dithiothreitol (DTT) in AMBIC and incubated for 20 min at 55°C. Iodoacetamide (7.5 μl of 100 mM) was added and the mixtures left for 20 min at 25°C in the dark. For protein digestion prior to MS, sequencing-grade trypsin (Promega) was added (0.5 μg) and the mixtures incubated for 20 hrs at 37°C. Reactions were terminated by addition of trifluoroacetic acid (0.5 μl) and the peptide mixtures were desalted by a modified Stage-Tip method as described previously [22]. LC-MS/MS analysis was performed using an Acclaim PepMap RSLC column of 50 cm × 75 μm inner diameter (Thermo Fisher Scientific) using a 2 hr acetonitrile gradient in 0.1% aqueous formic acid at a flow rate of 250 nl/min. Easy nLC-1000 was coupled to a Q Exactive mass spectrometer via an easy-spray source (Thermo Fisher Scientific). The Q Exactive was operated in data-dependent mode with survey scans acquired at a resolution of 75,000 at *m*/*z* 200 (transient time 256 ms). Up to ten of the most abundant isotope patterns with charge +2 or higher from the survey scan were selected with an isolation window of 3.0 *m*/*z* and fragmented by higher-energy collisional dissociation with normalized collision energies of 25. The maximum ion injection times for the survey scan and the MS/MS scans (acquired with a resolution of 17,500 at *m*/*z* 200) were 20 and 120 ms, respectively. The ion target value for MS was set to 10^6^ and for MS/MS to 10^5^, and the intensity threshold was set to 8.3 × 10^2^. Data were processed with MaxQuant (version 1.5) [43] and with PEAKS (version 7.5) [44] and peptides identified from the MS/MS spectra searched against combined reference human and herpesvirus type 1 (strain 17) proteomes [45]. For in silico digests of reference proteomes in MaxQuant the following peptide bond cleavages were allowed: arginine or lysine followed by any amino acid (a general setting referred to as trypsin/P). Samples in series were analysed together with the “match between runs” function enabled in the software. Up to two missed cleavages were allowed. The false discovery rate was set to 0.01 for peptides, proteins, and sites. Other parameters were used as pre-set in the software. In PEAKS, trypsin was selected as an in-silico enzyme and specific cleavages at both ends of the peptide were required and up to 3 missed cleavages allowed per peptide. The false discovery rate was set to 0.01 for peptides, and minimum of 1 unique peptide per protein requested. For HSV^AHA^ we unequivocally identified 77 Met→AHA substitution sites on 22 different viral proteins (Table S2, also indicated by red lettering in the analysis of total abundance, Fig 3c). By contrast, for HSV^wt^ tryptic peptides, only a single identification of an apparent Met→AHA substitution was returned. This provided a high level of certainty of correct peptide identification resulting from the very low false discovery rate.

### MS-based quantitative comparison of HSV proteins in HSV^AHA^ and HSV^Wt^ particles

For quantitative analysis of protein abundances in in HSV^AHA^ and HSV^wt^, samples of viruses were produced in parallel, and the raw files analysed and processed using a MaxQuant workflow. Two fixed modifications were selected, cysteine carbamidomethylation as well as a dummy-modification on methionine (Met→HPG). This second modification was introduced in order to remove methionine containing peptides from the analysis and thus exclude potential bias caused by underrepresentation of methionine containing peptides in raw files corresponding to HSV^AHA^ samples. Data were then further processed using Perseus software (version 1.5). Contaminants, reverse peptides, peptides identified by site only, and human hits were removed, and LFQ values converted to Log2. Protein LFQ intensity values corresponding to HSV^AHA^ and HSV^wt^ samples were plotted against each other to illustrate the comparative distribution of viral proteins in each virus sample (Figs 4C and S1).

### Detection of AHA-modified sites in HSV proteins

Purified samples of HSV^AHA^ and HSV^wt^ produced by either single-step or multi-round infection were processed as above and data analysed using PEAKS software. Cysteine carbamidomethylation was selected as fixed modification and methionine oxidation in addition to Met→AHA were selected as variable modifications. Spectra of AHA-peptides were manually inspected in PEAKS for unambiguous assignment of amino acid sequence and modification sites. HSV proteins were ordered based on the number of unique peptides identified and Met→AHA substitution sites were identified and listed (see Table S2).

### Estimation of AHA incorporation into HSV proteins by pulse-SILAC

RPE-1 cells (biological triplicates, 10^7^ cells/ test sample) were infected (moi 5) in serum free media. After 1 hr the inocula were removed and replaced with fresh media containing 2% serum. At 8.5 hpi, to deplete amino acid pools prior to dual SILAC and AHA labelling, this medium was removed and replaced with media lacking Met, Lys, and Arg (containing 2% dialysed serum). At 9 hpi the depletion medium was removed and replaced with media lacking Met, Lys, and Arg but supplemented with AHA (1 mM), ^13^C6,^15^N2-Lys and ^13^C6,^15^N4-Arg. At 24 hpi, the medium containing virus was harvested and extracellular virus particles clarified from cellular debris by low speed centrifugation (1,000 ×*g*, 10 min) and then pelleted and concentrated by high speed centrifugation (16,600 ×*g*, 1.5 h). The pellets was lysed in 2% SDS in PBS and sonicated briefly (2 × 5s) and further processed as in the MS sample preparation and workflow described above. Raw files were analysed with MaxQuant software. The fixed modifications cysteine carbamidomethylation, and dummy-modification on methionine (Met→HPG), were used as above, the latter to remove methionine containing peptides from the analysis and exclude potential bias caused by underrepresentation of methionine containing heavy peptides. Two variable modifications: R(+10Da) and K(+8Da) were also selected. Data were then further processed using Perseus software (version 1.5). Contaminants, reverse, identified by site only, and human hits were removed, and H/L values used to calculate efficiency of incorporation of AHA into HSV proteins according to the equation: % AHA = H/L ÷ (1 + H/L). The data are presented as a graph (Fig S2) and a table (Table S3).

## Acknowledgments

We are grateful to Guido Stoll for preliminary work in development of HSV labelling conditions and Todd Wisner for the generous provision of antibodies.

## Supporting Information

**S1 Fig. Comparative mass spectrometry analysis of viral protein content in HSV^AHA^ and HSV^wt^** Virus stocks were made during pulse-labelling with the A/M regime (or with Met alone) during single-step (panel a) or multi-step replication (panel b). Equivalent samples of viruses, normalised on the basis of infectivity were lysed, digested with trypsin, and analysed by LC-MS/MS. The results are plotted as the correlation between Log_2_ LFQ Intensities (approximating protein abundances) measured for proteins of HSV^AHA^ (Y-axis) and HSV^wt^ (x-axis). Proteins for which Met→AHA substitution sites were detected are highlighted in red. Representative proteins discussed in the text are circled. Further Data for Log_2_ LFQ Intensities are presented in Table S1.

**S2 Fig. Estimate of the degree of AHA incorporation into HSV proteins** (a) Schematic of labelling regime. Immediately after infection, cells were incubated in standard media. At 8.5 hpi, to deplete pools, this medium was removed and replaced with media lacking Met, Lys, and Arg. At 9 hpi the depletion medium was removed and replaced with media lacking Met, Lys, and Arg but supplemented with AHA, R^10^ and K^8^ (the latter two at the normal concentration for DMEM-F12 formulation). Virus was harvested after 24 hpi and virus particles purified and processed for MS. The percentage R^10^ and K^8^ incorporation was taken as a surrogate measure for the percentage AHA incorporation during the same labelling interval. (b) The data are illustrated where each vertical bar represents an individual, identified HSV protein and the Y-axis represents the % AHA incorporation into that protein during the labelling interval. ND means not detected. (c) The relative % incorporation for the population of virus proteins was binned into 10% ranges and the number of HSV proteins in each bin then plotted.

**S3 Fig. Quantitative analysis of HSV^AHA^ particles bound to cells by immunofluorescence and CuAAC ligation**. As for Fig 6, cells were infected with HSV^AHA^ and incubated +4°C then fixed immediately. Fig 6a represents the boxed section of the field shown here in panel a. Particles bound to cells at +4°C were detected by CuAAC ligation (green channel) versus detection by anti-VP5 capsids immunofluorescence (red channel). Panel a is a representative field of cells infected at +4°C which was quantitated using Image J as described in methods. Intensities for individual particles (ROIs) in each channel are shown in panel b with Y-axis the VP5 intensity and the X-axis AHA intensity. Each dot in the figure represents a particle ROI which is scored positive in a channel if it is 1 standard deviation above the mean background ROI for that channel (dotted lines). Particles that are positive for both signal are coloured orange, particles that are positive for AHA only are coloured green, and particles that are positive for VP5 only are coloured red.

**S4 Fig. Analysis of HSV^AHA^ and de novo VP5 synthesis.** Cells were infected with HSV^AHA^ as normal, shifted to 37°C for 6 hrs, fixed and the distribution of VP5 analysed. Arrows indicate cells with de novo synthesised nuclear VP5 observed at various levels. (b) Cells were infected in the presence of PAA (400 µg/ml) to block virus DNA replication and analysed 6 hpi for VP5 and AHA signals. The boxed area is shown as an inset with cytoplasmic VP5+ve capsids marked by arrows. These capsids are also AHA+ve. (c) An example of cells infrequently observed in the presence of PAA where large numbers of cytoplasmic particles could be observed. The inset shows that in such cases, virtually all capsids were also AHA+ve and thus represented incoming infecting particles.

**S1 Table. Quantitative analysis of the relative protein abundances in HSV^AHA^ and HSV^wt^.** HSV^AHA^ and HSV^wt^ stocks purified in parallel and equalised on the basis of infectious units, were subject to tryptic digestion and LC/MS as described in STAR methods. Raw files were processed using MaxQuant and Perseus software (version 1.5). The table gives LFQ values logarithmized (Log2) for three independent comparisons (preparations 1-3) of HSV^AHA^ and HSV^wt^. Preparation 1 was from stocks made by multi-step replication with the results shown graphically in Fig 3c while preparations 2 and 3 were from stocks made from single-step growth cycles. LFQ intensity values were plotted against each other to illustrate relative distribution of viral proteins (Fig 3C and S1 Fig).

**S2 Table. Identification of individual AHA-labelled protein species in HSV^AHA^** HSV^AHA^ was processed for MS as described in the text and materials and methods. For AHA detection, Met→AHA and methionine oxidation was selected as variable modifications. Spectra of AHA-peptides were manually inspected for unambiguous assignment of amino acid sequence and modification sites. The table lists HSV proteins where Met→AHA substitution sites were unambiguously identified and the corresponding individual amino acid number where Met has been replaced by AHA.

**S3 Table. Estimate of the degree of AHA incorporation into HSV proteins** Preparative stocks of HSV^AHA^ were made after pulse-labelling with AHA, K^+8^ and R^+10^ as described in the text and legend to S2a Fig. An estimate of the incorporation of AHA into specific proteins was calculated from the aggregate ratio of tryptic peptides for that protein that contain light or heavy isotopomers of Lys and Arg. The % incorporation for AHA was calculated from K^+8^ and R^+10^ as (H/L)/((H/L)+1) where H/L is the average intensity ratio of heavy to light peptides associated with each protein. Note that using our filter parameters for stringent identification of proteins in the biological replicates (samples 1-3), not all runs returned identified proteins. However when information was returned in at least one of the biological replicates this has been included in the summary table for approximation of AHA incorporation.

